# The functional brain favours segregated modular connectivity at old age unless targeted by neurodegeneration

**DOI:** 10.1101/463810

**Authors:** Luis R Peraza, John T O’Brien, Andrew Blamire, Marcus Kaiser, John-Paul Taylor

**Affiliations:** Institute of Neuroscience, Newcastle University, Campus for Ageing and Vitality, Newcastle upon Tyne, NE4 5PL, United Kingdom; Department of Psychiatry, University of Cambridge School of Medicine, Cambridge, CB2 0SP, United Kingdom; Institute of Cellular Medicine & Newcastle Magnetic Resonance Centre, Campus for Ageing and Vitality, Newcastle upon Tyne, NE4 5PL, United Kingdom; Interdisciplinary Computing and Complex BioSystems (ICOS) research group, School of Computing, Newcastle University, Newcastle upon Tyne, NE4 5TG, United Kingdom

**Keywords:** Resting state, Modularity, Connectome, fMRI, Lewy body disease, Alzheimer’s disease, Parkinson’s disease

## Abstract

Brain’s modular connectivity gives this organ resilience and adaptability. The ageing process alters the human brain, leading to modifications to its organised modularity and these changes are further accentuated by neurodegeneration, leading to disorganisation. To understand this further, we analysed modular variability—heterogeneity of modules—and modular dissociation—detachment from segregated connectivity—in two ageing cohorts and a mixed cohort of neurodegenerative diseases. Our results revealed that the brain follows a universal pattern of modular variability that matches highly active and metacognitive brain regions: the association cortices. We also proved that in ageing the brain moves towards a modular segregated structure despite presenting with increased modular heterogeneity—modules in older adults are not only segregated but their shape and size are more variable than in young adults. In the presence of neurodegeneration, the brain maintains its segregated connectivity globally but not locally; the modular brain shows patterns of differentiated pathology.

---

The brain can now be studied by a range of neuroimaging and electrophysiological technologies that allow us to further our understanding of its structure and function. By these recent developments, we know that brain function is dictated by complex interactions between neurons in the micro-, meso- and macroscales^1^, and that these are shaped in a functional network with specific properties and characteristics^2,3^. The functional brain network has been shown to be small-world, meaning that its structure reflects a balance between efficient communication and wiring cost^4,5^. The brain achieves this efficiency by creating groups of neurons densely connected among themselves but loosely communicated between the different groups; these are typically referred to as communities or modules^6^. It is hypothesized that the modularity of the brain is the result of evolution, where, in an always changing environment the brain developed a strategy to adapt subsystems rapidly without compromising the totality of its network^6^. In this regard, recent studies on brain dynamics have reported that modules constrain dynamic communication within their boundaries without affecting other modules^5,7,8^. This property also gives the brain a superior resilience against attacks either by disease or injury^4,9^.

Previous research in brain modularity have reported, consistently, several major modules such as the motor-sensory, visual and default-mode modules^10,11^. Although there is no agreement on their number, the majority of functional studies report that between three to ten modules are present in the brain^6,12^. This number depends on many factors and methodologic preferences such as number of brain regions (brain parcellation), the chosen brain atlases^13^, neuroimaging pre-processing pipelines^14^, connectivity measures (e.g. wavelet correlations), and treatment of connectivity weights^15^. Regardless of these differences, it is now agreed that brain modules are variable across time and between individuals, changing in shape and size depending on the cognitive task or no task at all, as is the case of resting state. A recent investigation by Bassett, et al. ^16^ in a task-based fMRI study found that after training, brain modules become more segregated and this was linked to learning and specialisation of regions. In the same fashion, Baum, et al. ^17^ reported that the modular brain becomes segregated during development, from childhood to adulthood, a finding reported as well by others^18^. From the perspective of diseases, alterations to brain’s modularity depend highly on the disease. In schizophrenia, for instance, there is a decrease in modularity suggesting leakage of information between modules^12^ while in the Lewy body diseases there is an increase in modularity suggesting segregated communities^19,20^. Certainly, the processes of ageing and neurodegeneration can alter brain communities; however how this is done and how the brain diverges from healthy ageing to a dementia state due to neurodegeneration is still not completely understood.

To address this issue, we studied functional modular changes by the ageing process within two public databases; the Enhanced Nathan Kline Institute Rockland Sample (NKI, N=297 participants)^21^ and the 1000 Functional Connectomes Project (TFC, N=359)^22^. Additionally, to test the effects of neurodegeneration on brain’s communities we also analysed a Newcastle University (NCL) database of neurodegenerative dementias^23–25^ which comprised an Alzheimer’s disease dementia group (ADD, N=42), and two Lewy body disease groups which included both dementia with Lewy bodies (DLB, N=38) and Parkinson’s disease dementia (PDD, N=17). These diseases are the most common cause of neurodegenerative dementia in older adults^26^ and represent a spectrum between a more cortical amyloid disease and subcortical alpha-synuclein disease that often concur in patients^27^. The NCL database also included age-matched healthy participants (N=34).

We investigated group modular variability (MV) which measures the heterogeneity of network communities across participants^28^. Additionally, we studied a proposed new measure we have named modular dissociation (MD). MD is defined as the community difference between two networks constructed by global and local thresholding of network connections. Global threshold is the standard and most used method for network construction while local thresholding is an alternative approach that favours segregation of modules by connecting the *k* nearest neighbours of a node, i.e. the strongest *k* connections. Hence, MD measures how close a globally thresholded network is from a nearest neighbour connectivity regime that favours modular segregation; thus high MD values indicate that the networks dissociate from a nearest neighbour connectivity regime.

We compared differences in MV and MD between young and older adults from the NKI and TFC cohorts and the deviation from healthy ageing to disease with the NCL cohort. This analysis was performed at optimal edge density (network cost) using a 451 regions of interest (ROI) functional atlas^29^. For validation purposes, group MV and MD were also estimated at 10% and 20% edge densities, and with three additional functional atlases; 100, 200, 247 ROI.

Our results show that the brain has a consistent pattern of high MV and MD across all studied cohorts which involves the higher association cortices and basal brain structures respectively, indicating a topographic and preserved universality to these measures/patterns. These patterns also demonstrated a consistent change as a result of healthy ageing, as observed in both independent ageing cohorts, where the brain moves towards a more segregated modular network structure that favours nearest neighbour connectivity despite presenting with increased modular heterogeneity, with the exception of the insulo-opercular cortex which showed higher values of dissociation and heterogeneity in older adults. However, in the presence of neurodegenerative dementia, the brain’s dissociation remains invariant—globally it stays in a segregated state. In contrast, it is heavily altered at the modular level with decreased MD in default mode and ventral-frontal modules and increased MD in the motor-sensory module. The present work represents an advancement in the understanding of the effects of ageing in the modular brain and the changes driven by neurodegenerative diseases on the ageing brain.

## Results

Participants from the ageing neuroimaging databases were selected and divided by age group: young adults (YA) between 20-40 years old (NKI =152 and TFC=257 participants) and older adults (OA) between 60-80 years old (NKI=146 and TFC=102 participants). The healthy participants within the NCL neuroimaging database were classified as an independent OA group and used as reference for comparisons with the three dementia groups. Demographics of all groups are given in Table S1.

### Length-vs-strength connectivity behaviour is not different among dementias

We examined edge distance vs edge strength (weight) behaviour within each group and performed comparisons between groups for each of the three cohorts in this study. Weights for this analysis belonged to the network connections that survived optimal edge density using local threshold network construction (Fig S2). We implemented a linear model to test for differences in the intercept and slope between two groups with connectivity strength as the dependent variable. In both ageing cohorts (TFC and NKI) a difference in the strength-vs-distance slope was found. Both older adult groups presented with a more negative slope compared with the YA groups indicating a faster decrease of connectivity strength as the Euclidean distance increases, which was significant at a p-value = 0.0002, Fig 2A-B. Differences in the intercept were found for the TFC cohort, with OAs showing a higher intercept compared with YAs (p-value < 0.0001) but no differences for this parameter were found for the NKI cohort. The covariate for sex was significant for the NKI group indicating differences in connectivity strength between males and females, p-value<0.0001. For the TFC cohort, sex was not a significant variable (p-value = 0.54).

**Fig 1.**
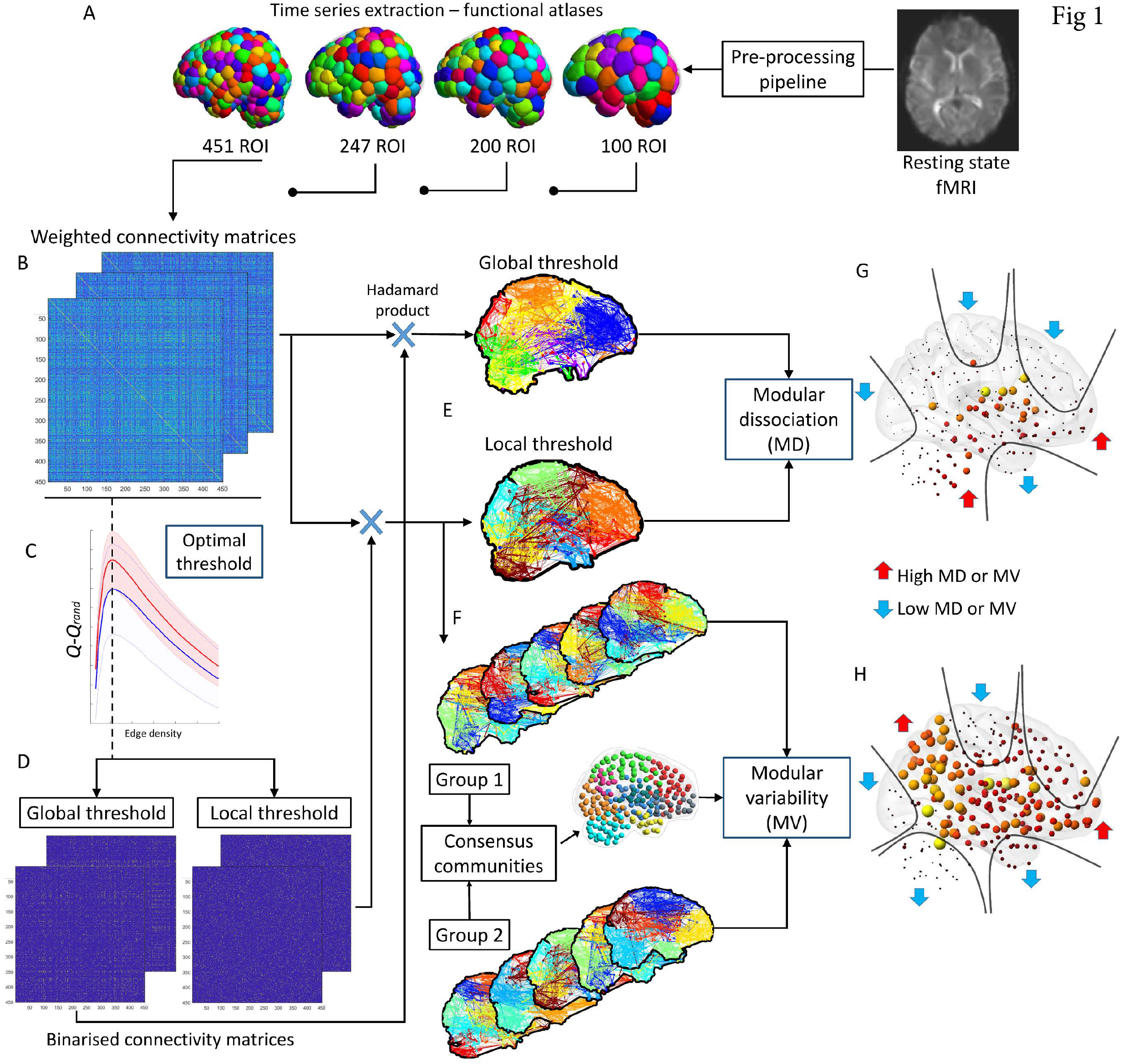
Methods for modular variability (MV) and modular dissociation (MD). A) Resting-state functional MRI pre-processing pipeline and time series extraction from functional atlases. B) Pearson correlation matrices. C) Optimal local threshold estimation; the network edge density at which *Q-Q_rand_* is maximum. D) Thresholded matrices by optimal density using local and global threshold network construction methods. E) Louvain’s community and modular dissociation (MD) estimation, see also Fig S1. F) Modular variability (MV) using consensus community. G) Group mean MD; subcortical regions and cerebellum showed in all groups high MD while motor-sensory, frontal, temporal pole and occipital cortex show low MD. H) Group mean MV; patterns of high and low MV were consistent across all groups. Motor-sensory, occipital, and temporal pole showed low MV while parietal, ventral frontal and insulo-opercular cortices showed high MV.

**Fig 2:**
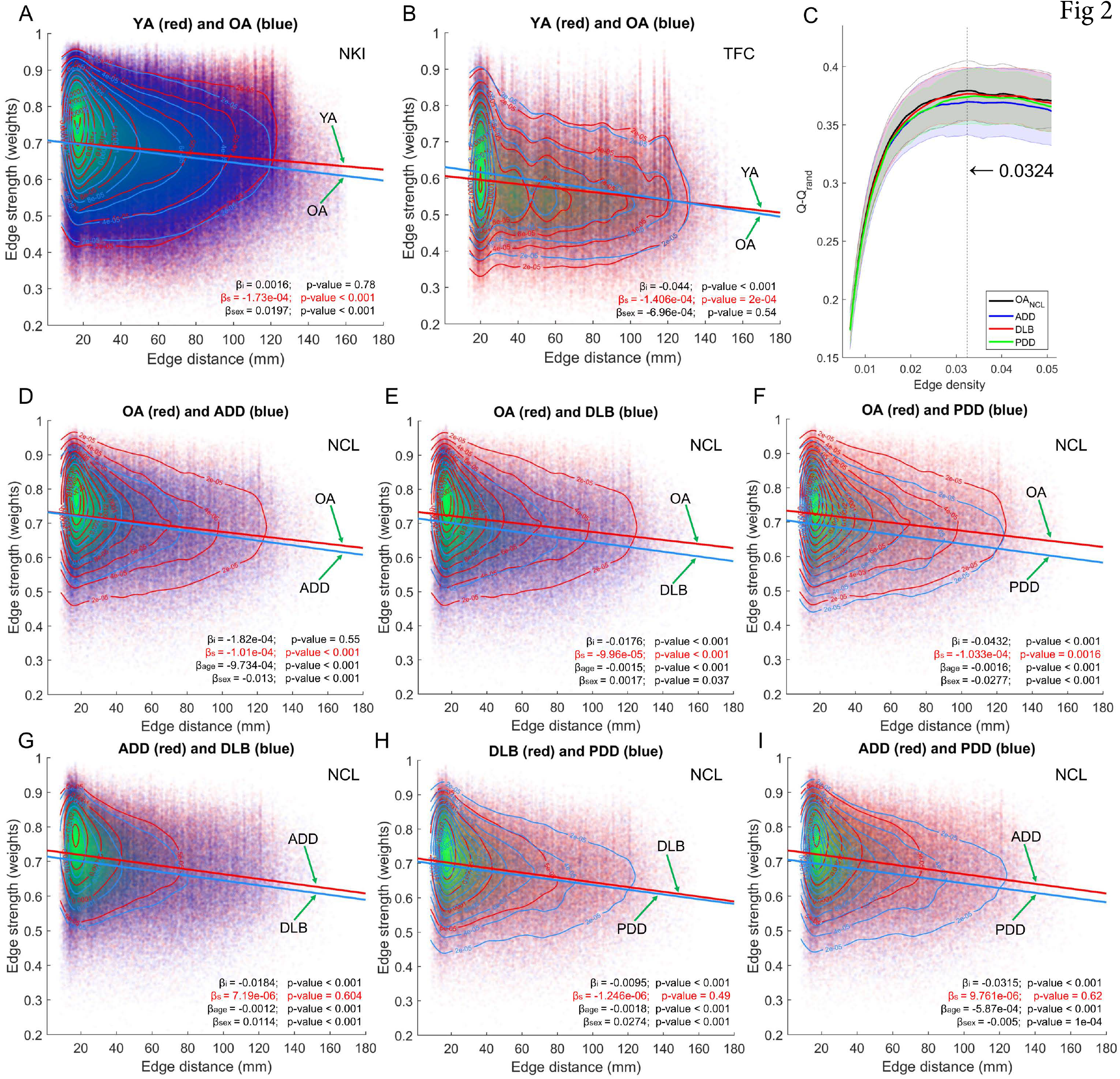
Network edge strength versus Euclidian edge distance. A) NKI; slope and intercept comparisons between edge strength and Euclidian distance profiles for the weighted thresholded connectivity matrices at optimal edge density, 3.8% with a 451-region of interest (ROI) atlas. B) TFC; slope and intercept comparisons for the weighted thresholded connectivity matrices at optimal edge density, 5.11% with a 177 ROI atlas. C) Optimal density estimation for the NCL cohort. Optimal edge density for all groups was reached at 3.24% with a 451 ROI atlas; see Fig S2 for the rest of cohorts. D-F) NCL; slope and intercept comparisons between edge strength and Euclidean edge distance profiles comparing OA vs ADD (D), OA vs DLB (E) and OA vs PDD (F). The three neurodegenerative dementias showed a significantly steeper slope when compared with OA. G-I) NCL; slope and intercept comparisons between dementias. The three dementia groups did not show differences in their slopes. Coefficients; β_i_ = intercept difference, β_s_= slope difference, β_age_= age, β_sex_= sex.

Fig 2C shows the estimation of *Q-Q_rand_* across a range of network edge densities for the NCL cohort, and which reached its maximum at 3.24% for the 451-ROI atlas, this edge density is referred as optimal density (see methods section); optimal density estimation for the other cohorts are shown in Fig S2. When comparing the distance-vs-strength profile for the NCL cohort, all dementias showed a significant negative slope that was steeper than the OA group (p-value < 0.0016). The ADD group showed no significant difference for the intercept when compared with OA (p-value = 0.55) whereas the DLB and PDD groups showed a significant lower intercept (p-value < 0.001). However, when comparing between dementias no significant slope differences were found, indicating that their strength-vs-distance connectivity profile was broadly similar, although their intercepts were different in all between-dementia comparisons (p-value < 0.001). Contrasting with the experiments for the ageing cohorts, age and sex were added as covariates of no interest. In all comparisons these covariates were significant, indicating that for the NCL cohort, age and sex influenced functional connectivity strength. Because of this, age and sex were added as covariates of no interest in all subsequent analyses.

### Modular variability is higher in association cortices than primary cortices

Due to the unbalanced group sizes in our study, we estimated MV using a bootstrapping approach where at each iteration an equal number of individuals from each group were randomly sampled without replacement and their community consensus estimated. Then, MV was computed as the variability between each selected participant and the consensus community at that iteration. These consensus communities were saved and a final meta-consensus estimated, and this is shown in Fig 3 at the top of each panel. For the NKI cohort, eight consensus modules were found (Fig 3A), seven modules in the TFC cohort (Fig 3B), and for the NCL a total of nine modules (Fig 3C). Several modules were consistent across cohorts. For instance, occipital and motor-sensory modules were found in the three cohorts. Additionally for the NKI and NCL, which had the same parcellation and data pre-processing pipeline, cerebellar, motor-sensory, and occipital modules were also consistent. There were some notable differences, however; for instance nodes within the basal brain were grouped with the insula cortex for the NKI cohort, while for the NCL cohort these same nodes were grouped with the cerebellum as a community.

**Fig 3.**
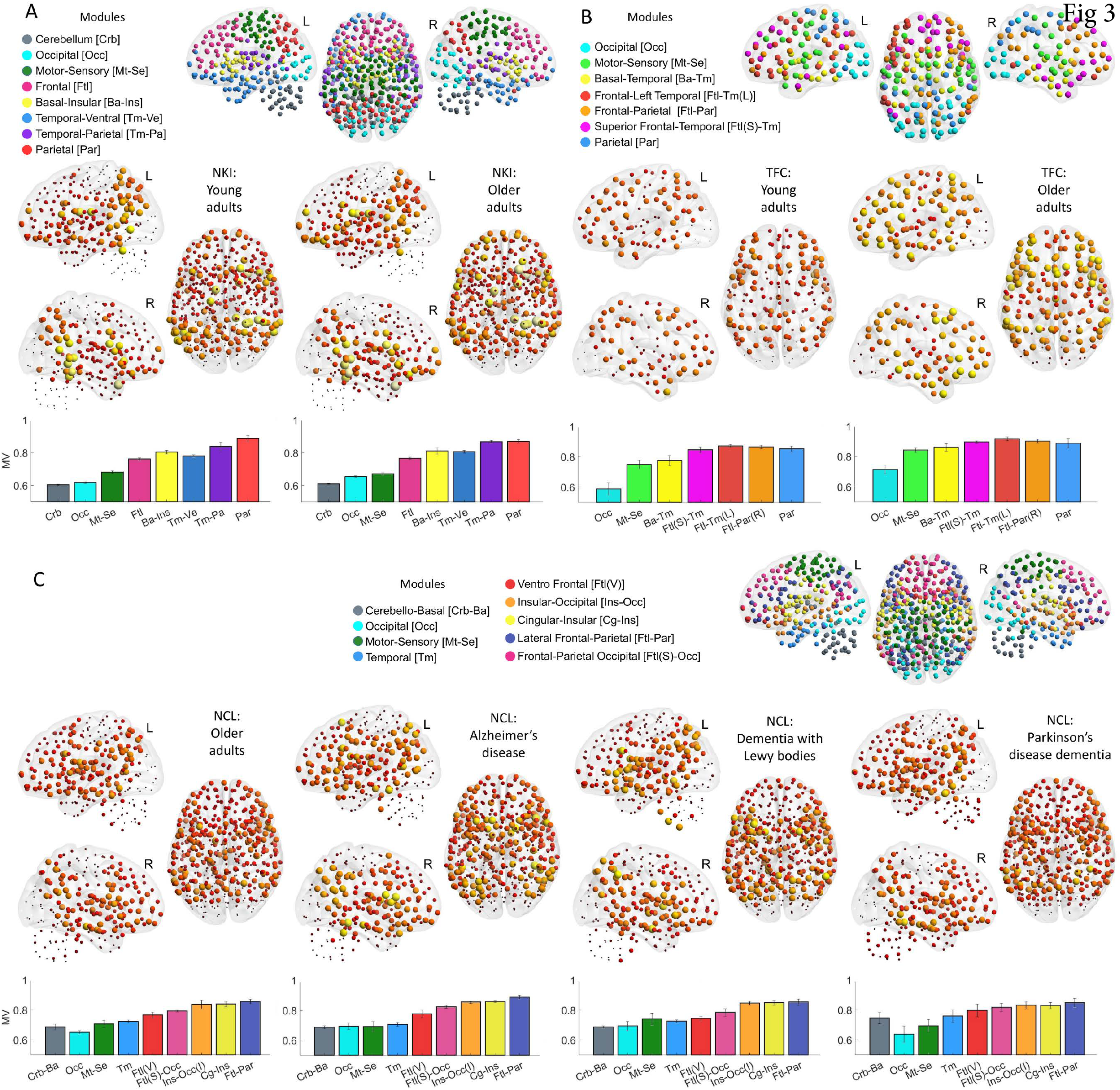
Modular variability (MV), consensus communities and group means. A) Nathan Kline Institute (NKI) consensus modularity and module definitions shown in coloured spheres (top). Group mean MV for young adult (YA) and older adult (OA) groups (middle). Mean MV per module shown as bar plots for the OA and YA groups (bottom); colours for each bar match the communities. B) Same as (A) but for the 1000 functional connectome (TFC) cohort. C) Same as (A) for the Newcastle (NCL) cohort. Left hemisphere (L), right hemisphere (R). Results presented here used the 451-ROI atlas for the NKI and NCL cohorts at optimal edge density; for results using other atlases see Fig S4.

We then studied group modular variability (MV) in our three study cohorts. Mean MV for all groups is shown in Fig 3A-C. All three independent cohorts and their subgroups showed a consistent pattern of MV across the brain. MV was higher at inferior frontal, parietal, insular cortices, inferior post and pre-central gyri as well as superior temporal gyri. Regions of low MV were the occipital, superior motor-sensory (or superior central gyri), temporal poles and the cerebellum (the latter for the NKI and NCL cohorts only). The superior frontal cortex also showed low MV but to a lesser extent compared with the occipital cortex. Even though the TFC cohort database did not have cerebellar connectivity, had a lower atlas resolution (177 ROI) and an independent pre-processing pipeline, this cohort also showed a similar MV pattern to the other two cohorts (Fig 3B).

The within module mean MV for all groups is shown as bar plots at the bottom of each panel in Fig 3. For all cohorts, the occipital and cerebellar modules showed the lowest MV values, followed by the motor-sensory modules. Modules with high mean MV were those with frontal, insular, and parietal aspects in their topological distribution; specifically parietal and insular cortices were the regions with the highest MV.

We also estimated modular dissociation (MD). In this case, MD can be estimated within each participant (i.e. bootstrapping was not necessary) and mean MD was computed directly. Similarly to MV, the pattern was consistent across the three cohorts although similarities were more evident between the NKI and NCL groups, and are shown in Fig 4A and C. For the TFC low MD is found in occipital, and pre and post-central gyri, while high MD is found in temporal poles and one node in the ventromedial prefrontal cortex, Fig 4B. NKI and NCL participant also showed low values of MD in occipital, and motor-sensory cortices while regions of high MD were located primarily in cerebellar, basal structures and insular cortices, Fig 4A and 4C.

**Fig 4.**
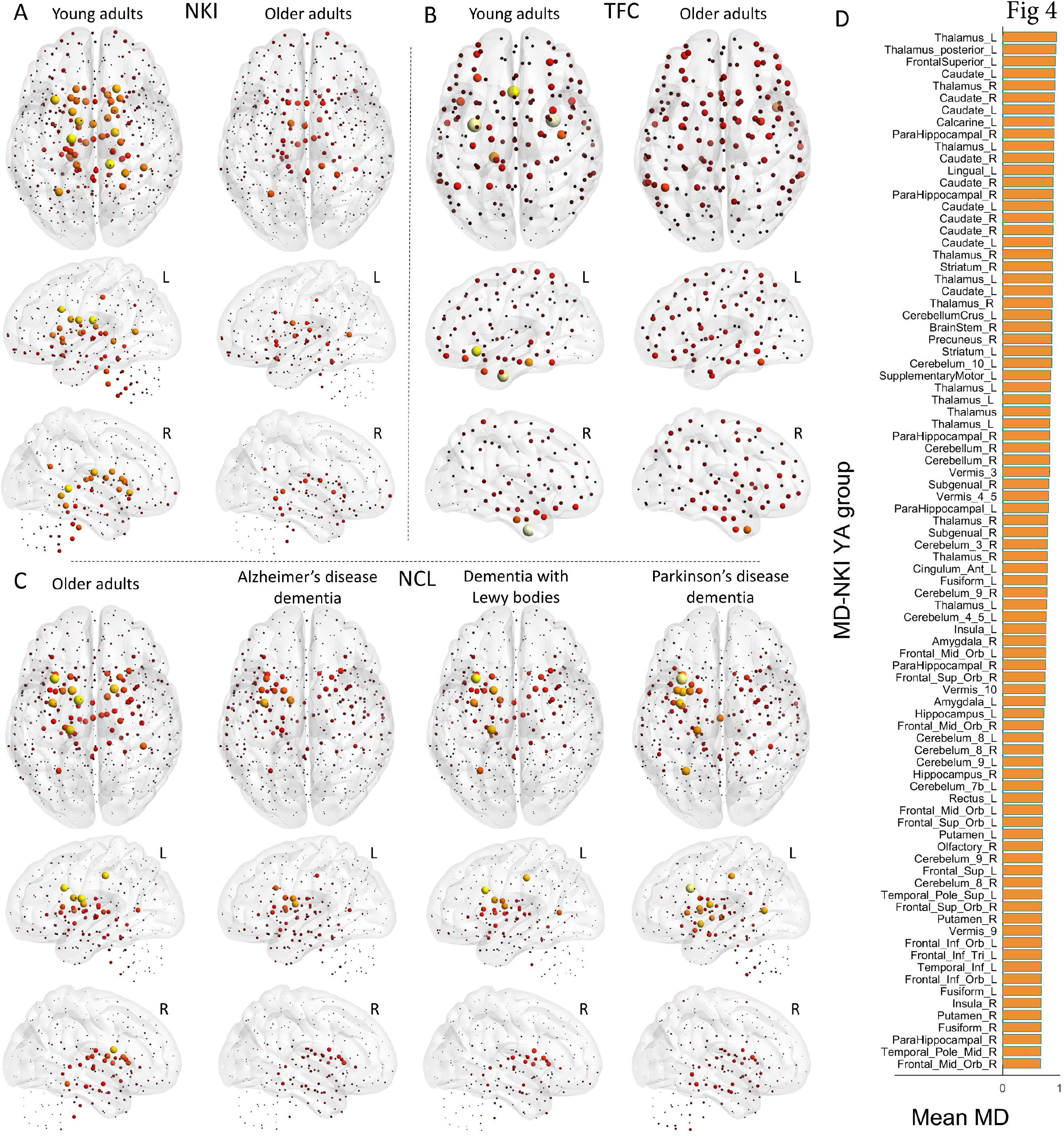
Mean modular dissociation (MD) for all study groups. A) Nathan Kline Institute (NKI) cohort MD values for the young adult (YA) and older adult (OA) groups. B) Same as (A) for the 1000 functional connectome (TFC) cohort. C) Same as (A) for the Newcastle (NCL) cohort. D) Strongest 20% of MD values within the NKI-YA group. Left hemisphere (L), right hemisphere (R). Results presented are from the 451-ROI atlas at optimal edge density; for results using other atlases and densities see Fig S5.

### Healthy ageing shows consistent patterns of increased modular variability and decreased modular dissociation

We performed between-group comparisons for MV and MD, and for the NCL cohort we were interested in comparisons between OA and the neurodegenerative dementias. All tests were corrected for age, sex and study by regressing out these covariates before nonparametric permutations for the between-group comparisons.

The TFC-OA group showed an overall increase of MV across the brain while the NKI-OA group showed regions of high and lower MV when compared with YA (Fig 5A-B). However, despite this difference both ageing groups showed a similar pattern of high MV within occipital, insular and ventral frontal cortices in OA compared with YA. For the NKI groups, OA showed lower MV within parietal, motor-sensory, cerebellum and part of the superior frontal cortices (Fig 5A).

**Fig 5.**
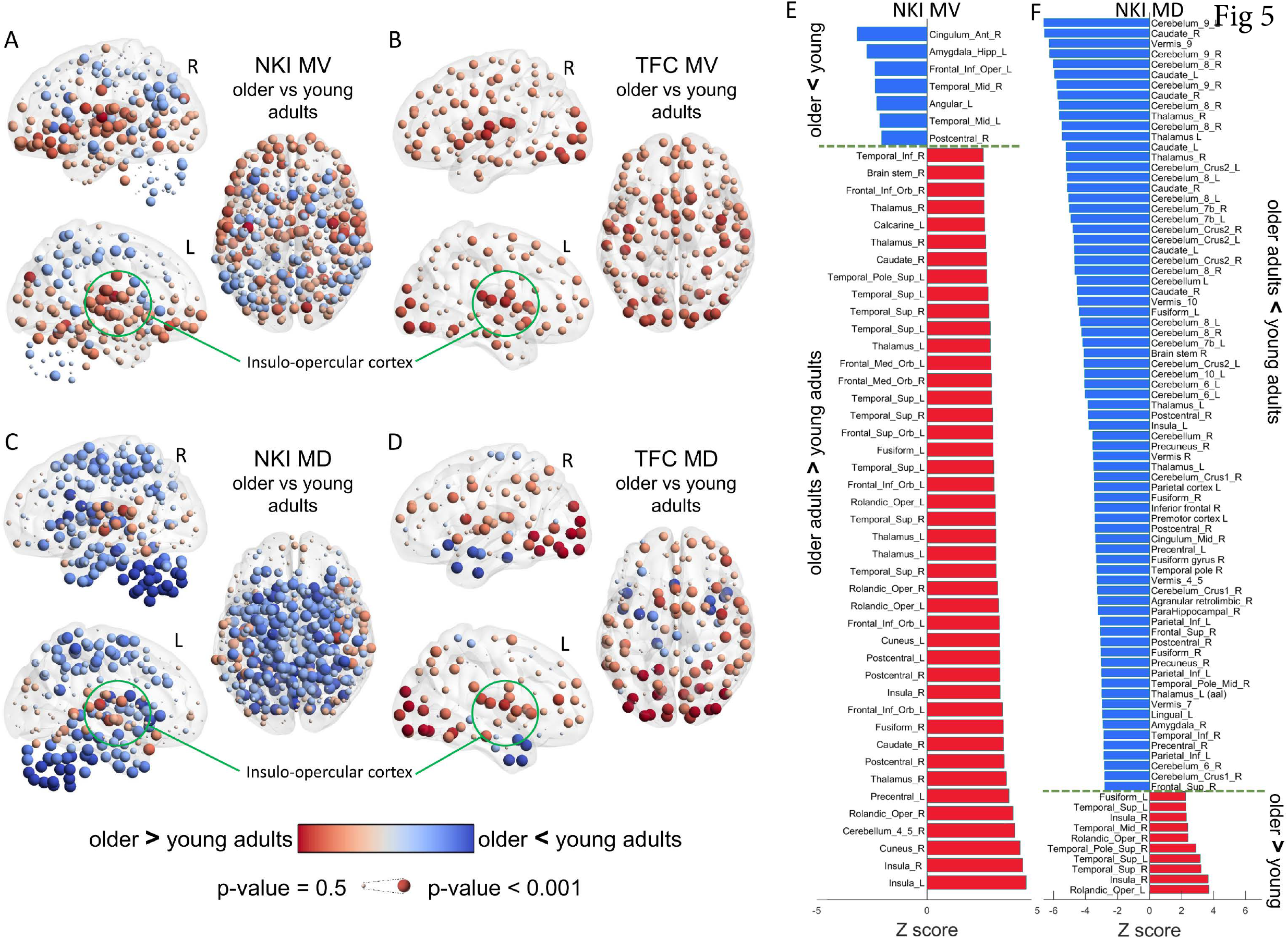
Age effects of modular variability (MV) and modular dissociation (MD). A) Age differences in MV within the Nathan Kline Institute (NKI) cohort, older adults (OA) compared with young adults (YA). B) Age differences in MV within The 1000 functional connectome (TFC) cohort OA compared with YA. C) Age differences in MD within the NKI database, OA vs YA. D) Age differences in MD within the TFC cohort, OA vs YA. E) Significant brain regions from the NKI-MV comparison; corrected for multiple comparisons at p-value<0.05. F) Significant brain regions for the NKI-MD comparison; corrected at p-value<0.05. Differences assessed with non-parametric permutations (5000) after regressing covariates of no interest. Regions’ names are given using the Automatic Anatomical Labelling (AAL).

The pattern for MD for both ageing cohorts, NKI and TFC, was also similar. Again, insular areas showed significantly higher MD in older adults while cerebellum, temporal and motor-sensory regions showed lower MD in OA compared with YA (Fig 5C). For the TFC cohort, in particular, the occipital cortex showed significant higher MD in OA compared with YA (Fig 5D); this trend also existed within the NKI cohort, but it was not significant after corrections for multiple comparisons. Between-group comparisons for both ageing groups survived FDR correction for multiple comparisons at a p-value < 0.05. Corrected results for the NKI comparisons for MV and MD are shown in Fig 5E and Fig 5F respectively.

### Neurodegenerative dementias show differentiated patterns of modular variability and dissociation

When comparing dementia patients with OA within the NCL cohort, the ADD group showed on average higher MV within the occipital cortex while lower MV was found in the rest of the cortex, Fig 6A. The DLB group also showed higher MV in the occipital cortex as well as in the motor-sensory cortex. Surprisingly, the DLB group showed regions that were statistically different, but the direction of this difference (higher or lower than OA) changed between iterations during the bootstrapped meta-analysis, resulting in low differences (shown with pale grey colour) but significant and represented by large spheres. These regions were located primarily within the insular and ventral frontal cortices, Fig 6A middle panel. For the PDD group, differences were not significant for MV (uncorrected) compared with OA, with only a trend of high MV within the cerebellum, Fig 6A left panel.

**Fig 6.**
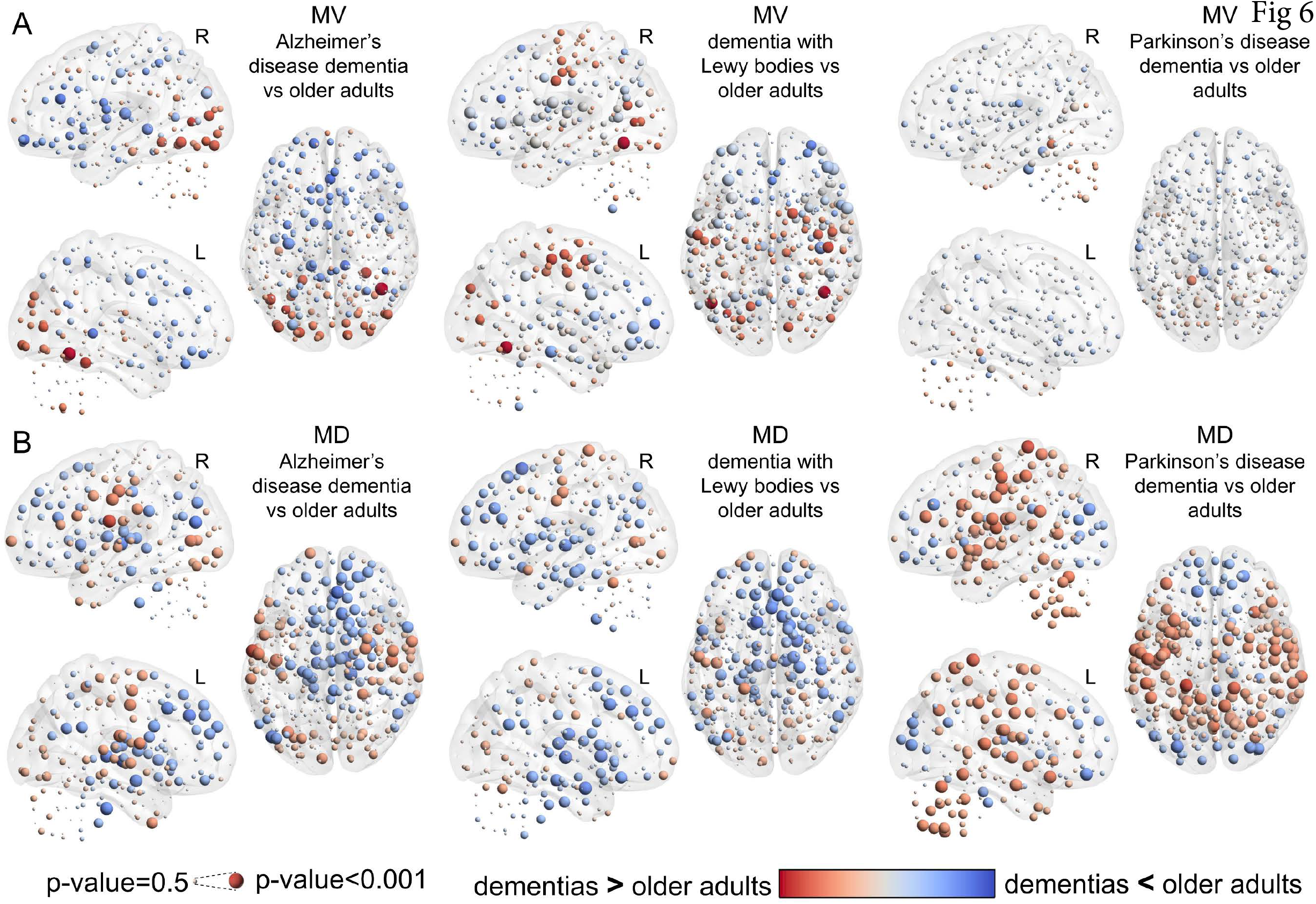
Neurodegenerative dementia effects of modular variability (MV) and modular dissociation (MD). A) MV differences between older adults (OA) and dementia groups: Alzheimer’s disease dementia (ADD), dementia with Lewy bodies (DLB), and Parkinson’s disease dementia (PDD). B) MD differences between OA and dementias. Results assessed with non-parametric permutations (5000) after regressing out covariates of no interest. Results for MV-PDD were not significant (uncorrected) while the rest of the comparisons showed significant uncorrected results. Only results for MD-PDD survived correction for multiple comparisons at a p-value < 0.05.

MD was also compared between dementias and OA, where the ADD and DLB groups showed a similar pattern of MD with higher values in occipital and motor-sensory cortices and lower values within frontal, ventral frontal, precuneal and basal brain regions. PDD, on the contrary, showed a differentiated pattern of MD compared with the other two dementias, with lower MD at occipital cortex and high MD within cerebellar, insulo-opercular and motor-sensory cortices, Fig 6B.

### The functional brain moves towards a segregated modular structure with ageing whereas neurodegeneration alters this segregation locally

We further decided to explore MD differences between young adults and the neurodegenerative dementia groups. This was possible by using the healthy OA groups within the NKI and NCL cohorts as reference groups; MD values were analysed relative to the OA groups to create an MD_/OA_ index (see methods section). Fig 7A shows MD values from both OA groups. There was a high agreement between both OA groups regarding MD; Pearson r = 0.8, p-value=1.17e-102, R^2^=0.64. Fig 7B shows the global MD across all nodes; here MD was significantly higher in YA compared with OA globally while this measure did not change significantly in our exemplar dementia groups indicating that globally MD is not affected by neurodegeneration; i.e. MD_/OA_ was significantly different from zero in YA but not in the dementia groups. However, when we analysed MD changes relative to OA for each of the identified communities, higher YA MD was present in six of the nine modules, as shown in blue in Fig 7C; ventral frontal, fronto-parietal (or default mode), temporal, motor-sensory, cerebello-basal, and lateral fronto-parietal modules. For the modules with a frontal component, the MD change was on average negative in the dementia groups (Fig S3 C, D and H), indicating lower MD compared with OA. For the temporal module, the ADD group showed a positive change although this was significantly lower than YA MD changes (Fig S3-E), and for the cerebello-basal module the PDD group showed on average a positive MD change but this was significantly lower than the YA MD change. For the motor-sensory module, all groups showed a positive change relative to OA but this change in dementias was significantly lower than YA (Fig 7C). From these results, it is worth noting that for the occipital module, the PDD group showed a significant negative change of MD which contrasted with the positive MD change in ADD and DLB groups (Fig S3-G).

**Fig 7.**
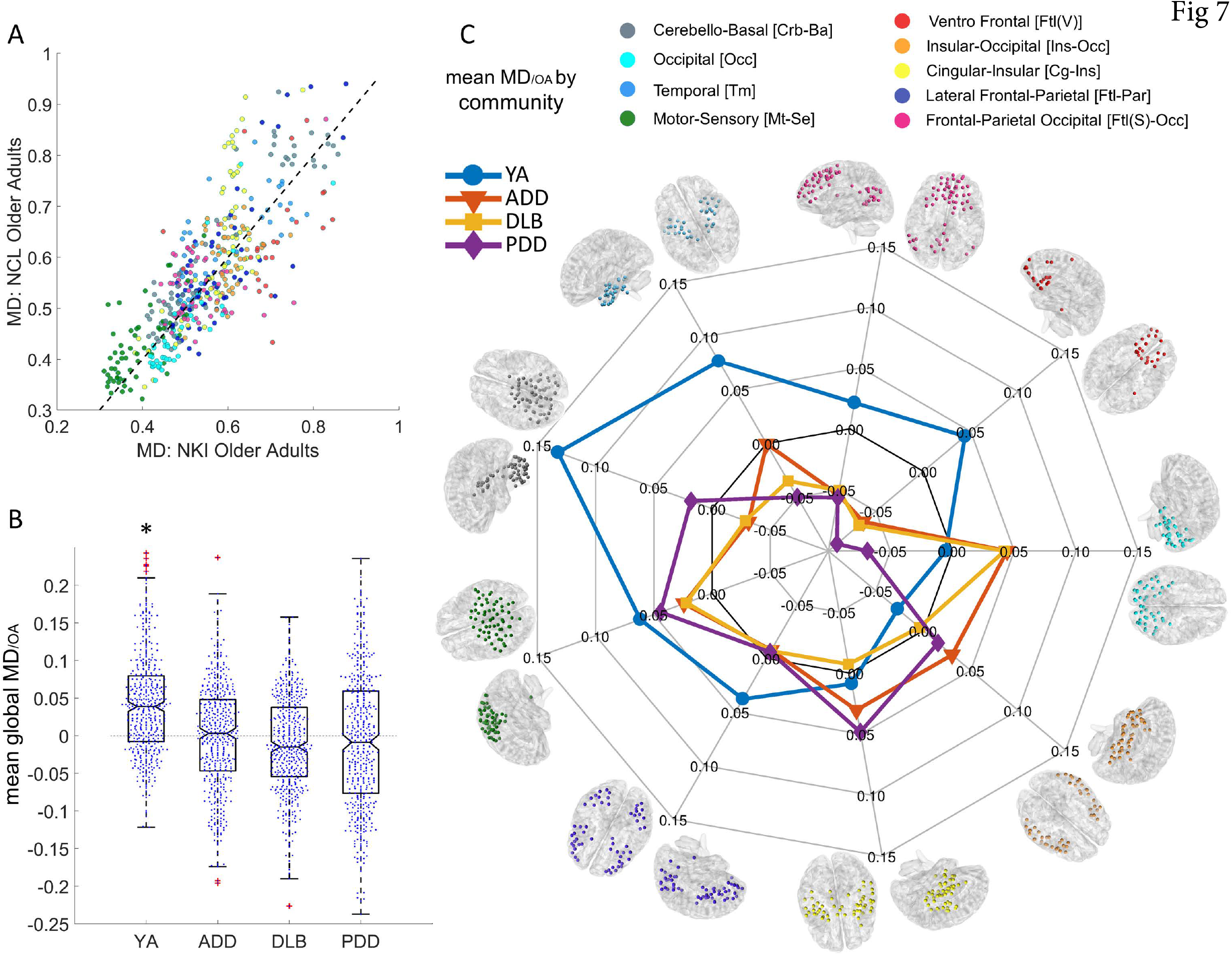
Modular dissociation relative to older adults, MD_/OA_; changes from healthy ageing to dementia. A) Scatter plot with MD values from the OA groups within the Nathan Kline Institute (NKI) and the Newcastle University (NCL) cohorts; colours are shown according to communities. Both OA groups were correlated; Pearson r=0.8, p-value = 1.17e-102, R^2^=0.64. B) Global MD_/OA_ for the NKI-young adult (YA) group and the NCL groups; Alzheimer’s disease dementia (ADD), dementia with Lewy bodies (DLB), and Parkinson’s disease dementia (PDD). *Statistically different from zero, Wilcoxon signed rank test: p-value = 1.183e-29. C) Spider plot for Mean MD_/OA_ by community. Ageing drives a significant decrease of MD shown by the high scores in YA group (blue line) and mainly for the cerebello-basal community, suggesting that the ageing brain moves towards a local connectivity regime or segregation. Neurodegenerative dementia did not change overall brain’s MD_/OA,_ but alterations are present at different communities. See Fig S3 for non-referenced results of MD values and box plots.

## Discussion

Our research study showed consistent patterns of group modular variability (MV) and modular dissociation (MD) across the three independent neuroimaging cohorts and which suggest pattern universality for these two characteristics. Our main findings are that in the ageing brain, its modules move towards a connectivity regime that favours segregation while brain modules become more heterogeneous in older adults compare with young adults. This shift towards a segregated modular connectivity is not affected by neurodegeneration globally. However, at the modular level, there are marked changes where modules showed increased and decreased modular dissociation depending on the type of neurodegenerative disease.

### Modular variability is high in association cortices and low in primary cortices

Our method of modular variability measures the variability in community assignment of a node when compared against the group’s consensus community. Hence, MV measures modular heterogeneity across participants. This measure was observed to be high within association cortices: the posterior and anterior association cortices, which span the parietal and frontal lobes. The limbic association cortex which spans the temporal pole did not demonstrate high MV.

In contrast, the insular cortices showed high MV in all groups. The brain regions showing high MV are known brain areas considered of high demand; i.e. that these engage with multiple modules or brain systems and are involved in multiple cognitive tasks^7^. Indeed, the regions of high cognitive activity identified by Bertolero, et al. ^7^ are remarkably similar to our pattern of high MV across participants. Hence, our MV pattern may be capturing the different interactions between nodes of high demand that connect with a great variety of regions and modules.

Brain regions with low MV were the motor-sensory, occipital cortex and cerebellum. The motor-sensory and occipital cortices are known primary cortices that receive information from thalamic connections^30^. These network modules are also the most consistently reported in the literature indicating that these are highly integrated in the functional brain and that nodes within these modules interact less often with other brain regions^31^. The cerebellum was found here to be a module with low MV, this structure is highly connected to the basal ganglia^32^ and it is topologically isolated from the cortex, with the dentate nucleus as its main route to the basal brain^33^ and which may drive its consistent modular integrity.

A hypothesis for the differences in modular variability between the association and primary cortices is that during the neurogenesis, the association cortex develops later in humans allowing flexibility of function and behaviour^34^, whereas primary cortices develop earlier with strong connections from the basal brain, which may also control their cytoarchitecture and functionality^35^. The high modular variability observed in particular within the insular regions and operculum seems to indicate high demand. The insula in the human brain is characterised by the presence of von Economo neurons, which are fast communication circuits within the brain^36^. Additionally, it has been reported that the insula is involved in multiple brain functions, regulating salience stimuli and activity between brain systems such as the attention and the default mode systems^37^.

### Mean modular dissociation follows a similar pattern to modular variability

The patterns for MD and MV showed low values within the motor-sensory and occipital cortices/modules indicating that at the group level these modules are very homogeneous across participants and are densely connected within their respective communities i.e. segregated by nearest neighbour connectivity.

The MD pattern showed remarkably high values within basal regions in all groups. Because of the definition of MD, this index measures the difference between global and local threshold construction methods. This difference is mainly driven by the weakest connections introduced by the local thresholds and which change the topography of communities. These connections are not weak in correlation given that these belong to the strongest nearest neighbours for each node within the participant’s connectivity matrix. However, the higher MD values within the basal region indicate that nodes within this region have a high tendency to disconnect or join other modules, i.e. dissociate. In this regard, the basal brain and specifically the thalamus presents with weak and sparse connections to the cortex^30^, even though these connections can control the global functional dynamics of the whole cortex^38^.

The structure of weak connections in the brain has been a matter of intense research. These connections are found to be non-random and their function is thought to relate to network path length shortening^39^. Also, previous research has found that the weak long-distance connections contribute to the brain’s complexity^5^ and that their strength decreases with cortical distance^40^, a finding that we also confirmed here (in Fig 2). Previous investigations in modular connectivity have used the strongest connections using either static^10^ or dynamic^16^ approaches. It is highly plausible that because of the weak and dynamic nature of the basal connections, these previous studies were not sensitive to basal connectivity, even though these are highly targeted by a wide range of brain diseases and conditions^41^.

### Ageing patterns of variability and dissociation are universal

The comparisons between ageing groups, OA vs YA, from both cohorts showed similar patterns of MV and MD suggesting that the ageing process follows a consistent path of alterations. In addition, our results demonstrate that generally, the brains of OA become more segregated with lower dissociation values compared with YA. The motor-sensory, temporal and cerebellum which are regions of low MD, have this metric further decreased in OA while regions within insular and occipital cortices show increased MD. It is hypothesised that modular segregation is a result of functional specialisation^17^. In this regard, Baum, et al. ^17^ found using structural networks (from diffusion tensor imaging) that modules segregate from childhood to adulthood, which agrees with our results using functional connectivity and further extrapolates this conjecture to older adults. Baum et al. also reported that the opercular cortex does not follow segregation with age which agrees with our result in functional networks. The high MV and MD shown by the OA group in the insulo-opercular cortex compared with YA were observed in both NKI and TFC cohorts indicating a universal characteristic.

The occipital cortex also showed higher MD and MV in OAs from both ageing cohorts, although this characteristic was more prominent in the TFC cohort. The occipital cortex is one of the most resilient regions to ageing; cortical thickness in this region does not significantly decreases with ageing^42^ and shows minimal thinning/atrophy in ADD^43^. However, this cannot explain the higher MD and MV observed in OA because cortical thickness within the insular cortex, contrary to the occipital cortex, is significantly reduced by ageing^42^. At this point, our only explanation is that both regions are of higher demand in the old age brain.

When we compared MV and MD between dementias and older adults, only results for the comparison PDD vs OA survived correction for multiple comparisons. This also indicated that for ADD and DLB, our two nodal indices remained fairly invariant to the presence of these neurodegenerative dementias. However, a pattern was observed in ADD and DLB; these dementias showed a trend of higher MD in the occipital and motor-sensory cortices. This contrasted with the NKI and TFC cohorts where OA showed lower MD for these regions compared with YA, which suggest that MD in motor-sensory cortex partially returns to YA’s levels in the presence of neurodegeneration and that this region departs from a modular segregation regime that is a characteristic of ageing.

PDD showed higher MD in motor-sensory and insular cortices as well as in the cerebellum; only occipital and frontal cortices showed a trend that favours segregation of modules. These differences may be related to the neuropathology of the Lewy body diseases affecting motor and cognitive brain functions; however, more research on larger cohorts will be needed to bring more light into this.

### Neurodegenerative dementia does not further alter brain’s segregation globally, but it does at the modular level

The global mean MD across the cortex did not change significantly with the presence of dementia, contrary to normal ageing which showed lower MD in OA compared with YA. However, and despite this global invariance, at the modular level there were significant changes in all dementia groups. Six of the nine modules showed lower relative MD than the YA group. This first indicated that with ageing, the brain is more segregated, with nodes closer in relative strength to their communities. However the effects of dementia vary, for some modules the tendency is of higher or lower MD depending on the type of dementia but in many of these cases this was significantly different from zero, indicating change. For instance, the negative relative MD in all dementias for the ventral-frontal module (Fig 7C in red) shows that during ageing, nodes within this module follow a connectivity regime that favours modular segregation, and that in the presence of neurodegeneration, nodes follow this tendency even further (negative MD_/OA_). The contrary occurred with the motor-sensory module, where the positive relative MD in all dementias shows that this module is partially ‘restored’ to young adult levels of dissociation. The pattern of MD_/OA_ between ADD and DLB is very similar, confirming the similarities between these two diseases. Surprisingly, the PDD group showed a completely different MD_/OA_ pattern compared with DLB, even though previous investigations have suggested functional similarities between DLB and PDD^25,44^.

### The importance of the weak connections

Previous investigations have focussed their attention on the strongest connections. This is certainly sensible; the strongest connections are less influenced by noise and are less likely to be of artefactual origin. However, recent investigations have shown the importance of the weak or slightly weaker connectivity within the brain. In a resting state fMRI investigation linking brain connectivity with intelligence quotients (IQs) from participants, Santarnecchi, et al. ^45^ found that the 60-80% range of the weakest end of the edge-weight distribution, correlated with IQ, and this correlation was higher and more significant than for the strongest connections at the 1-20% range. In our investigation, we performed a test to observe where in the connectivity matrix the weak edges were added by the local thresholding method (Fig S6). Most of the added edges by the local threshold construction method were intra-modular edges, i.e. these edges made communities concise. On the contrary, edges added by global threshold were inter-modular, or in other words these aimed to fuse modules; especially the motor-sensory and temporo-parietal modules, Fig S6-C and D. Additionally, the number of intra-modular edges (by their mean node degree from binarised edges) added by local thresholding correlated positively with global MD, proving their influence to the MD index.

The network construction method by local thresholding showed interesting properties in our investigation. Using this method, the modularity statistic Q was not different between dementia groups and OA, and the same can be seen for the NKI database of OA and YA, Fig S1. Also for both databases, Q variance is lower when using local thresholding method. Another observation is that the mean number of modules estimated by local threshold is the same for all groups within each of the databases (NCL=9, TFC=7 and NKI=8 modules at their respective optimal edge densities). This is because the local thresholding method is based on the *k*-nearest neighbour graph, which favours local connectivity by connecting the closest nodes and inducing a more segregated network. This was confirmed by analysing the edge Euclidean distance distribution for the connections introduced by the local threshold network construction regime, where close range connections are highly introduced. Also, the long-range weaker connections although less probable than short range, are more probable than in a global threshold regime (Fig S7). In conclusion, the weaker connections added by local thresholding, and which connected nearest neighbours, did not aim to communicate between modules but to consolidate modules, even in the presence of disease, in our case, neurodegenerative dementia.

Fig 8 shows an explanation for the introduction of short-range weaker connections by the local thresholds in neurodegeneration. Currently, it is accepted that neurodegenerative diseases target specific brain systems or modules^46^; neurodegenerative diseases affect connectivity within these specific systems and their short-range connections^47^. Despite being affected by neurodegeneration, these local links are still connecting the nearest neighbours within their respective communities and can survive a local connectivity regime (Fig 8C). The finding that modularity statistic Q (Fig S2) and the number of modules where not different between dementias and OA using local thresholds suggest this.

**Fig 8.**
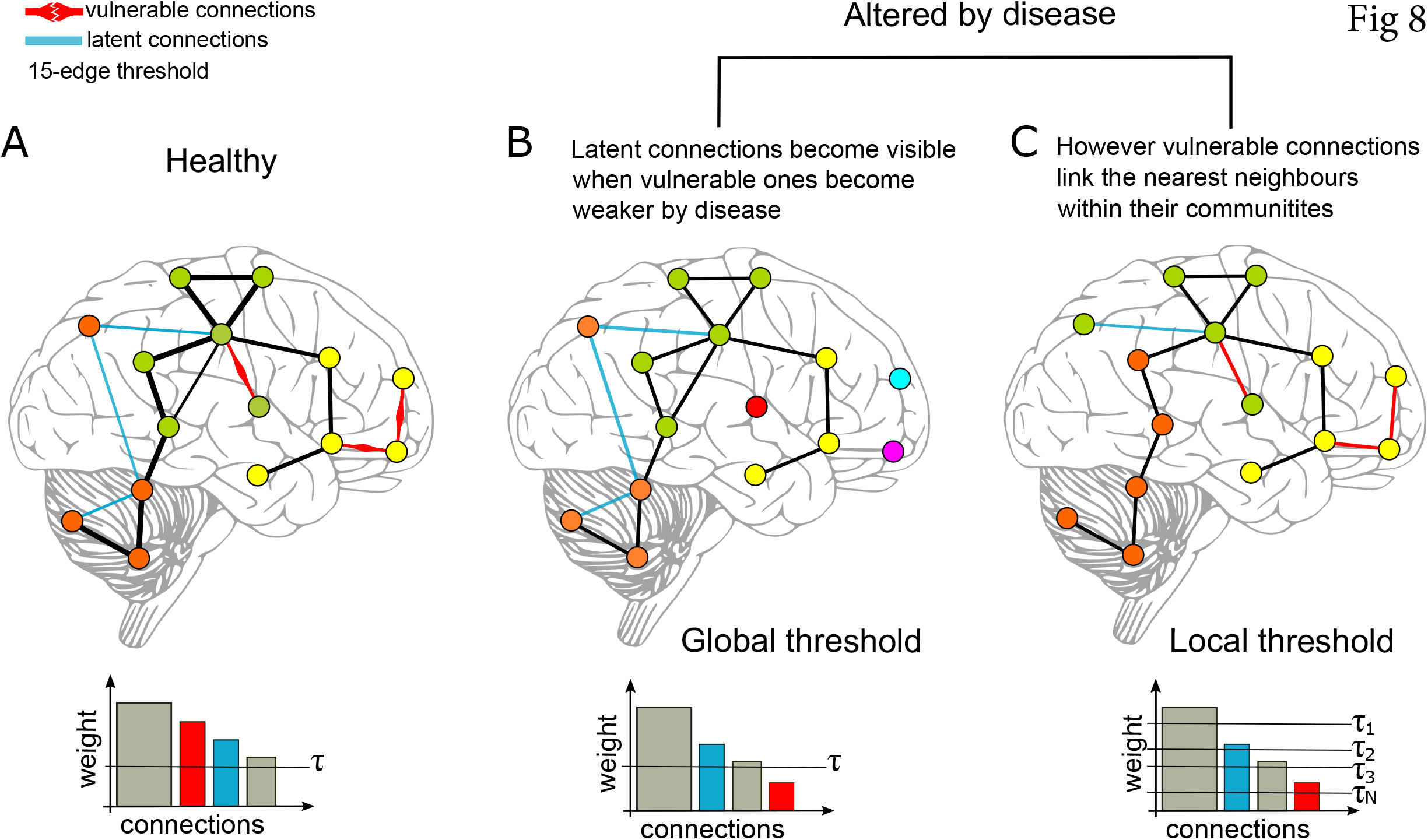
Disease effects on local and global thresholding. A) A toy model of healthy brain connectivity with “known” connections including vulnerable edges which can be targeted by disease, and latent connections which are weaker than vulnerable connections. B) When the model is tarjeted by disease, the vulnerable connections become weaker and do not survive a global threshold. C) Although weakened by disease, the vulnerable connections still connect the nearest neighbour nodes within their communities, and these connections are restored or made visible by local thresholds.

In conclusion, high modular variability or heterogeneity of modules exists within the association cortices which are known regions of high cognitive demand. Our results showed that this is a universal characteristic of the brain, whose changes are shown to be consistent in healthy ageing. Similarly, modular dissociation was consistently high at basal brain regions, and this brain characteristic decreases with healthy ageing, indicating that the brain moves towards a connectivity regime that favours segregation of modules, except for the insulo-opercular cortices which depart from this regime. In contrast, modular dissociation is not affected by neurodegenerative dementia globally but at the modular level. The brain is constantly learning and also gets experienced. The ability of the brain to dissociate modules in youth may be a reflection of continuous learning and the need for neuronal modules to communicate and share resources. As the brain gets older, these interactions get either impaired by ageing or less necessary due to experience and the brain moves towards a more segregated network structure. Despite this, brain modules in older adults are more heterogeneous across participants suggesting that the brain within individuals may follow different strategies or paths during the ageing process.

## Methods

### Experimental model and participant details

#### Newcastle participants and recruitment

Two independent neuroimaging databases were combined from two clinical studies^23–25^. Participants in these studies were recruited within the north-east of England and patients with neurodegenerative dementia were contacted through old-age psychiatry and neurology services in Newcastle area (United Kingdom). A total of 42 dementia with Lewy bodies (DLB, N=16 in study 1 and N=24 in study 2), 44 Alzheimer’s disease dementia (ADD, N=16 in study 1 and N=28 in study 2) and 17 Parkinson’s disease dementia (PDD, N=17 in study 2 only) patients were recruited. Additionally, 34 age-matched healthy control participants (N=16 in study 1 and N=18 in study 2) were recruited as a comparison group. All patients were diagnosed by two experienced clinicians according to the clinical criteria for these diseases: The dementia with Lewy bodies consensus criteria^48,49^, the diagnostic criteria for PDD^50^, and the National Institute on Aging-Alzheimer’s Association criteria for AD^51^. For both studies, approval was granted by the Newcastle Ethics Committee and all participants gave informed consent.

#### Enhanced Nathan Kline Institute (NKI) Rockland Sample

Participants from the NKI-Rockland sample^21^ were selected and neuroimaging datasets downloaded. Selection criteria comprised participants with resting-state functional magnetic resonance imaging (fMRI) who were between 20-40 years old for the young adult group (YA, N=151) and participants between 60-80 years old who comprised the older adult group (OA, N=151). Details about the recruitment of participants for this cohort can be found in Nooner, et al. ^21^.

#### The 1000 functional connectome project

Functional connectivity matrices from the 1000 functional connectome^22^ were downloaded from the USC Multimodal Connectivity Database (UMCD, http://umcd.humanconnectomeproject.org/)^52^. This connectivity repository comprises resting state connectivity matrices (Pearson correlations) from nine independent studies with a wide age range. A young adult group (YA, N=257 participants between 20-40 years old) and an older adult group (OA, N=102 participants between 60-80 years old) were selected and functional connectivity matrices downloaded.

### Neuroimaging procedures

#### Neuroimaging acquisition from Newcastle participants

Neuroimaging was acquired in both studies with the same scanner. Structural magnetic resonance imaging (MRI) was recorded with a 3T Philips Intera Achieva Scanner. Acquisition protocol used a magnetisation prepared rapid gradient echo (MPRAGE) sequence, sagittal acquisition, echo time 4.6 ms, repetition time 8.3 ms, inversion time 1250 ms, flip angle 8°, SENSE factor = 2, and an in-plane field of view of 256×256 mm^2^ with a slice thickness of 1.2 mm yielding a voxel size of 0.93×0.93×1.2 mm^3^ for study 1. For study 2, an in-plane field of view of 240×240 mm^2^ with a slice thickness of 1.0 mm yielding a voxel size of 1.0×1.0×1.0 mm^3^ was used. For functional resting-state neuroimaging, both studies used the same recording protocol: gradient echo echo-planar imaging sequence with 25 contiguous axial slices, 128 volumes, anterior-posterior acquisition, in-plane resolution = 2.0 × 2.0 mm^2^, slice thickness = 6 mm, repetition time = 3000 ms, echo time = 40 ms, and field of view = 260 × 260 mm^2^.

Neuroimaging protocols for the NKI and TFC databases can be found in Nooner, et al. ^21^ and Biswal, et al. ^22^ respectively and references therein.

#### Neuroimaging pre-processing

Neuroimages from the NCL and NKI databases were pre-processed with the same pipeline and blinded to the study groups. Non-brain tissue was stripped with the brain extraction tool (BET) from the FMRIB Software Library (FSL version 5.0)^53^ and results visually inspected to check if brain structures were complete and isolated. Resting-state functional MRI (rs-fMRI) were then processed. For the NKI resting state neuroimages, the first five volumes were deleted because the time series were not steady. All rs-fMRIs were motion corrected using the FMRIB’s Linear Image Registration Tool (FLIRT) without spatial smoothing. Then, the six-movement variables from FLIRT and the average time series from the bilateral ventricle were regressed out from all resting state images. In order to further correct for movement and other artifacts, the rs-fMRI time series were despiked with the BrainWavelet Toolbox^54^; parameters were adjusted to despike approximately 0.6% of the times series. Structural and functional images were then coregistered using FEAT, and nonlinear coregistration to MNI152 space was implemented with the FMRIB’s Nonlinear Image Registration Tool (FNIRT) within FSL. Then, fMRI time series were high-pass filtered with a 150-second filter using FSL-FEAT. Finally, all functional images were transformed to a 4×4×4 mm^3^ resolution, and a 6-mm full-width half maxima (FWHM) spatial smoothing was applied to all volumes.

Four functional brain parcellations or atlases were estimated with the pyClusterROI toolbox^29^: <=100, <=200, <=250, and <=500 regions of interest (ROI) were requested to the algorithm. For this, an independent dataset of healthy older adults (N = 29 participants recruited within the north-east of England and who had neuroimages recorded using the same scanner as the NCL cohort), were pre-processed with the same pipeline as the NCL and NKI databases. The functional brain from the independent group was parcellated with the pyClusterROI software using a grey matter mask from the Harvard-Oxford atlas (from FSL) at 0% threshold, and which included the cerebellum. Details for the independent older adult database can be found in Peraza, et al. ^55^ The functional atlases estimated and used in this study comprised 100, 200, 247 and 451 ROI. The average time series within each ROI was extracted from all rs-fMRIs, and Pearson correlation matrices were computed for each participant from the NCL and NKI databases. Main manuscript shows results using the 451- ROI atlas while results using the other functional atlases are given in the supplementary material.

The TFC matrices comprised Pearson correlations between 177 ROIs from a functional atlas^29^ and had no cerebellar structures; only the neocortex and subcortical structures were included. Further details about the pre-processing pipeline used to estimate these matrices can be found in Brown, et al. ^52^

After neuroimaging pre-processing, two AD and four DLB participants from the NCL and five OA participants from the NKI cohort were excluded due to coregistration inaccuracies.

#### Community and modularity statistic estimation

At this stage, there are three sets of connectivity matrices with Pearson correlation coefficients: NCL, NKI and the TFC cohorts. Currently, there is no consensus about the treatment of negative values from Pearson correlation matrices for the definition of network connections, and several approaches have been proposed such as deletion of all negative correlations^56^, transformation to an all-positive scale^57^ or compute the absolute value of the correlation coefficients |*r*|^19^. Here we decided to take the absolute value of the connectivity matrices as we did in our previous study^19^ because we were interested in the connectivity strength between regions rather than in the direction of these correlations. Furthermore, previous evidence from functional connectivity studies have proved the biological origin^58,59^ and importance of negative correlations in the brain^60^, further confirming our decision of keeping this connectivity information.

Networks were represented as binary undirected connectivity matrices. For this, the weighted matrices need to be thresholded and binarised. The thresholds were selected by edge density also known as network connectivity cost or proportional thresholding^15^. Network cost is defined as *2T*/(*N*^2^ − *N*) ×100%, where *T* is the total number of edges that survived the threshold and *N* the number of nodes. Three network edge densities were studied: optimal edge density (defined below), 10% and 20% of the strongest connections.

Modularity statistics were estimated using weighted thresholded matrices. For this, the Hadamard product between the binarised matrix and the original weighted matrix was computed, and used for community structure and modularity statistic estimation using the Louvain’s algorithm function from the Brain Connectivity Toolbox (BCT)^2^ in Matlab (Mathworks Inc, R2017a). In all Louvain community estimations, the algorithm was run 1000 times and the optimal community structure with the highest modularity statistic *Q* was saved for further analysis.

#### Network construction methods

When thresholding connectivity matrices by edge density, there are two possibilities on how this threshold is applied in order to build the binarised adjacency matrix: local^12^ and global threshold^61^. In the global threshold approach, a single threshold τ is applied to the entire weighted connectivity matrix selecting a percentage (cost) of the strongest edges; connectivity weights below a threshold τ are set to zero and above τ are set to one. Local threshold on the other hand, imposes a minimum node degree in all nodes within the network. Local thresholding is based on the *k* nearest neighbour graph (*K*-NNG) where every node is connected to its *k* nearest (strongest) neighbours^62^. Local thresholding works by searching the closest lower *K*-NNG, closest to the desired edge density, and the method adds extra edges to each node until the desired cost or the next *K*-NNG is reached^12^. Software to construct networks by local thresholding is provided as supplementary material.

As consequence, local threshold allows the inclusion of weaker connections in other to maintain the rule of *k* nearest neighbours and favours integration of communities and segregation of modules within the network. Indeed, a previous investigation noted that constructing networks by local threshold enhances the identification of communities^63^. Nevertheless, the weaker connections added by the local threshold construction regime would not be present when using a global threshold approach, and the structure of both binarised networks will differ mainly at these weaker connections^15^. A toy example for the differences between local and global threshold construction methods is shown in Fig S1 and a neurophysiological interpretation is shown in Fig 8.

#### Optimal cohort edge density selection

The optimal cohort edge density was selected as the density that maximised the difference between the participant’s network modularity and an equivalent random network modularity statistics, *Q*-*Q_rand_* ^6^. Modularity of the equivalent random network was estimated as the average modularity statistic of 40 randomised thresholded weighted matrices using the BCT function randmio_und_connected.m. A range of densities for each participant (from the NCL, NKI, and TFC cohorts) and atlas parcellations (100, 200, 247 and 451 ROIs) were explored for optimal edge density. From 0.3% to 25% (with 100 equally spaced increments) for the 100-ROI parcellation, from 0.2% to 10% (with 73 equally spaced increments) for the 200 and 247-ROI parcellations, and from 0.08% to 5% densities (with 51 equally spaced increments) for the 451-ROI parcellation. For optimal edge density discovery, we opted for Newman’s algorithm^64^ instead of Louvain’s because the former is faster to compute, and at this step we were not interested in the community structure *per se* but the edge density where the modularity statistic is maximally higher when compared with an equivalent random network. Once the optimal density was found, Louvain’s algorithm was implemented instead.

#### Connection length and strength comparisons

In order to test for connectivity differences in Euclidian distance and strength profile between groups, all weighted connections from the locally thresholded matrices at optimal edge density (see previous section on optimal edge density) were investigated with multiple linear regression models. For each group comparison, a linear model to test for differences in the intercept and slope of the distance-vs-strength regression lines between the two groups was implemented. This model was defined as *strength* ~ *β_1_·distance* + *β_2_·group* + *β_3_·group*\**distance* + *β_4_·age* + *β_5_·sex* + *β_6_·study* +*1*, where the coefficients *β_2_* and *β_3_* account for group differences in intercept and slope respectively. However, because of the correlation values within the weighted connectivity matrices |*r_ij_*| did not follow a normal distribution, bootstrapped linear models were implemented instead^65^. At each iteration, 20% of the total distance-strength connection population was randomly sampled without replacement together with their respective covariates for age, sex and study/site (if applicable). For each group comparison, 10000 bootstrapping iterations were run, linear model coefficients estimated and the coefficient histograms analysed to assess if these coefficients, which represent differences in slope and intercept, were significantly different from zero (p-value < 0.05).

#### Modular variability and modular dissociation

Modularity is defined as the ability of the system to be decomposed into communities or subsystems^11^. Functional modules are highly variable between and within individuals^28^ and previous investigations have proposed novel methods to study network modules, and their variability or heterogeneity in shape across time^16^ and even neuroimaging modalities^66^. Here, we are proposing two approaches to study modular variability (MV) between groups and between network construction methods (local and global threshold) which we have named modular dissociation (MD).

MV measures the module affiliation variability of a respective node *s*^67^. Given a node *s* and two modular partitions obtained from two participants or methods *i,j*, its variability *MV_s_*(*i*, *j*) is defined as^28^:

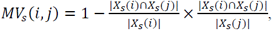

where |*X_s_*(*i*) ∩ *X_s_*(*j*)| represents the number of nodes in common between communities *X_s_*(*i*) and *X_s_(j)*, and |*X_s_*(*i*)| represents the number of nodes within community *X_s_*(*i*), where node *s* belongs to in the network *i* and similarly for |*X_s_*(*j*)|. MV ranges from 0 to 1, where 1 means no overlap between communities and 0 means no change in the node set comprising both communities^67^.

MD measures the variability between community definitions of two networks from the same participant but constructed using two thresholding regimes, i.e. modular variability between two networks constructed by local and global thresholding methods. Computation of MD is the same as for MV, but between network construction methods and consequently, MD will measure community allegiance between a global threshold which allows the strongest connections within the weighted matrix and a local threshold regime which allows weaker connections but favours community segregation by connecting the nearest neighbours. Also, note that both networks will have by definition the same edge density or cost. A toy example of MD estimation is shown in Fig S1, and a schematic figure showing the estimation procedure is shown in Fig 1.

### Quantification and statistical analysis

MV was assessed between study groups using locally thresholded networks^12^. This was decided because the modularity *Q* is not statistically different between groups in the NCL and NKI cohorts using this network construction method. Furthermore, the optimal edge density is reached at higher connectivity costs and led to higher *Q* statistics compared with global threshold for the three databases in this study, see Fig S2.

The analyses in this investigation are static analyses, which differs from previous investigations where MV has been assessed as the community change across time^10^. In order to test MV between two groups, we estimated the group’s community consensus^68^ with the Network Community Toolbox (NCT, http://commdetect.weebly.com/). This approach is more accurate than estimating communities from the group’s mean weighted connectivity matrix. The consensus community is estimated from the optimal communities obtained with Louvain’s algorithm, which was applied to all participants from the two groups. Then, MV for each participant and node is estimated as the difference between the participant’s Louvain community and the two-group consensus community. However, since none of our groups is perfectly balanced, i.e. we do not have an equal number of participants in all groups, there is a high risk that the consensus community is biased towards the largest group. To resolve this, we estimated MV with a bootstrapping approach. Having two groups of size *A* and *B* were *B* < *A*, we randomly choose without replacement *B* participants from each group, their consensus community estimated and nodal MV computed for each participant. This procedure was repeated 500 times. At each of these bootstrapped iterations, nodal MV differences between groups were assessed with nonparametric permutations (5000 permutations) after regressing out age, sex and study covariates. For the NKI and TFC cohorts, the age covariate was demeaned within groups only in order to correct for age variances without losing the group effect (YA vs OA). For the TFC cohort, study covariates could only be regressed within OA and within YA groups since neuroimaging acquisition for these two groups were recorded at different sites^22,52^. For the NCL cohort we added the study covariate, and the NKI analysis did not use a covariate for different studies. All results from the 500 iterations were averaged to obtain meta-results: p-value_meta_ and meta Z-score differences Z_meta_. For the Z scores, the mean and variance from the nonparametric permutations were used for normalisation.

From the previous analyses, a meta-consensus community was estimated for each of the study cohorts. For this, a community consensus was estimated again from all 500 consensus communities that resulted from the bootstrapping operation.

In order to estimate between-group differences for MD, the bootstrapping approach is not necessary because MD is computed within participants. Differences between groups for MD were assessed directly with nonparametric permutations (5000 permutations) after regressing out age, sex and study covariates in a similar fashion as MV. MV and MD results were corrected for multiple testing using false discovery rate (FDR) with the Benjamini and Hochberg procedure in Matlab; mafdr.m function at p-value < 0.05 for significance level.

Additionally, mean modular comparisons were also investigated relative to the OA groups; MD_/OA_. For this, nodal MD values from a group of interest were concatenated with values from the OA group (from the same cohort) to create a two-group MD vector *[G_1_ G_2_]* per node, where *G_1_* represents the OA values. Age, sex and study covariates were regressed out and a residual node vector *[R_1_ R_2_]* obtained. From this regression, mean nodal MD was estimated and values referenced to the mean OA residuals; *[R_1_ R_2_]-E(R_1_)*, where *E* stands for expected value. This latter step allows to compare MD between different cohorts, e.g. NCL and NKI, and define MD_/OA_ = MD_R2_-MD_R1_, where MDR1 are the regressed nodal MD values of the OA groups and MD_R2_ are the nodal MD values of the second group of interest. Differences between OA and the groups of interest were assessed with nonparametric two-sided rank sum tests; here nodes were treated as independent observations and results corrected for multiple tests.

#### Visualisation of results

Sphere brains were plotted using the BrainNet Viewer Toolbox^69^. For both MD and MV, results were nonlinearly mapped for visualisation, and for visualisation only, with an exponential function: *MD′* = 100*^MD^* and *MV′* = 100*^MV^*, where prime indicates visualised. This transformation mapped results from a 0-to-1 scale to a nonlinear 1-to-100 scale. A similar approach was followed for the visualisation of the between-group comparison maps: *pvalue′* = 100^(1−*pvalue*)^. Colours were implemented with diverging colour maps from blue to red colours^70^.

Identification of significant structures in MNI space was implemented with the brain anatomical database within the xjView toolbox (http://www.alivelearn.net/xjview/), and the cuixufindstructure.m function in Matlab.

## Data and code availability

Matlab code for MV, MD estimation and network construction by local thresholding are available at https://github.com/LuisPerazaRo/modulardissociation and by contacting the corresponding author and senior author. The Enhanced Nathan Kline Institute (NKI) Rockland Sample is available at http://fcon_1000.projects.nitrc.org/indi/enhanced/index.html and the 1000 Functional Connectome (TFC) resting state matrices are publicly available at http://umcd.humanconnectomeproject.org.

## Acknowledgements

The research was funded by a grant from Alzheimer’s Research UK in partnership with Hidden Hearing (ARUK-PPG2016A-2) to Luis R Peraza, Marcus Kaiser, and John-Paul Taylor, and supported by the National Institute of Health Research (NIHR) Biomedical Research Centre (BRC) at Newcastle University. The study —participant recruitment and data collection at Newcastle University— was funded by an intermediate clinical Wellcome Trust Fellowship (WT088441MA) to John-Paul Taylor.

## References

1 Bassett, D. S. & Bullmore, E. T. Human brain networks in health and disease. Current opinion in neurology 22, 340–347, doi:10.1097/WCO.0b013e32832d93dd (2009).

2 Rubinov, M. & Sporns, O. Complex network measures of brain connectivity: uses and interpretations. NeuroImage 52, 1059–1069, doi:10.1016/j.neuroimage.2009.10.003 (2010).

3 Kaiser, M. A tutorial in connectome analysis: topological and spatial features of brain networks. NeuroImage 57, 892–907, doi:10.1016/j.neuroimage.2011.05.025 (2011).

4 Achard, S., Salvador, R., Whitcher, B., Suckling, J. & Bullmore, E. A resilient, low-frequency, small-world human brain functional network with highly connected association cortical hubs. The Journal of neuroscience: the official journal of the Society for Neuroscience 26, 63–72, doi:10.1523/JNEUROSCI.3874-05.2006 (2006).

5 Gallos, L. K., Makse, H. A. & Sigman, M. A small world of weak ties provides optimal global integration of self-similar modules in functional brain networks. Proceedings of the National Academy of Sciences of the United States of America 109, 2825–2830, doi:10.1073/pnas.1106612109 (2012).

6 Sporns, O. & Betzel, R. F. Modular Brain Networks. Annual review of psychology 67, 613–640, doi:10.1146/annurev-psych-122414-033634 (2016).

7 Bertolero, M. A., Yeo, B. T. & D’Esposito, M. The modular and integrative functional architecture of the human brain. Proceedings of the National Academy of Sciences of the United States of America 112, E6798–6807, doi:10.1073/pnas.1510619112 (2015).

8 Kaiser, M., Görner, M. & Hilgetag, C. C. Criticality of spreading dynamics in hierarchical cluster networks without inhibition. New Journal of Physics 9, 110 (2007).

9 Kaiser, M., Martin, R., Andras, P. & Young, M. P. Simulation of robustness against lesions of cortical networks. The European journal of neuroscience 25, 3185–3192, doi:10.1111/j.1460-9568.2007.05574.x (2007).

10 He, Y. et al. Uncovering intrinsic modular organization of spontaneous brain activity in humans. PloS one 4, e5226, doi:10.1371/journal.pone.0005226 (2009).

11 Meunier, D., Lambiotte, R., Fornito, A., Ersche, K. D. & Bullmore, E. T. Hierarchical Modularity in Human Brain Functional Networks. Frontiers in neuroinformatics 3, 37, doi:10.3389/neuro.11.037.2009 (2009).

12 Alexander-Bloch, A. F. et al. Disrupted Modularity and Local Connectivity of Brain Functional Networks in Childhood-Onset Schizophrenia. Frontiers in Systems Neuroscience 4, 147, doi:10.3389/fnsys.2010.00147 (2010).

13 van Wijk, B. C. M., Stam, C. J. & Daffertshofer, A. Comparing Brain Networks of Different Size and Connectivity Density Using Graph Theory. PloS one 5, e13701, doi:10.1371/journal.pone.0013701 (2010).

14 De Vico Fallani, F., Richiardi, J., Chavez, M. & Achard, S. Graph analysis of functional brain networks: practical issues in translational neuroscience. Philosophical transactions of the Royal Society of London. Series B, Biological sciences 369, doi:10.1098/rstb.2013.0521 (2014).

15 van den Heuvel, M. P. et al. Proportional thresholding in resting-state fMRI functional connectivity networks and consequences for patient-control connectome studies: Issues and recommendations. NeuroImage 152, 437–449, doi:10.1016/j.neuroimage.2017.02.005 (2017).

16 Bassett, D. S. et al. Dynamic reconfiguration of human brain networks during learning. Proceedings of the National Academy of Sciences 108, 7641–7646, doi:10.1073/pnas.1018985108 (2011).

17 Baum, G. L. et al. Modular Segregation of Structural Brain Networks Supports the Development of Executive Function in Youth. Current biology: CB 27, 1561–1572.e1568, doi:10.1016/j.cub.2017.04.051 (2017).

18 Meunier, D., Achard, S., Morcom, A. & Bullmore, E. Age-related changes in modular organization of human brain functional networks. NeuroImage 44, 715–723, doi:10.1016/j.neuroimage.2008.09.062 (2009).

19 Peraza, L. R., Taylor, J. P. & Kaiser, M. Divergent brain functional network alterations in dementia with Lewy bodies and Alzheimer’s disease. Neurobiology of aging 36, 2458–2467, doi:10.1016/j.neurobiolaging.2015.05.015 (2015).

20 Ko, J. H., Spetsieris, P. G. & Eidelberg, D. Network Structure and Function in Parkinson’s Disease. Cerebral cortex, 1–15, doi:10.1093/cercor/bhx267 (2017).

21 Nooner, K. B. et al. The NKI-Rockland Sample: A Model for Accelerating the Pace of Discovery Science in Psychiatry. Frontiers in neuroscience 6, 152, doi:10.3389/fnins.2012.00152 (2012).

22 Biswal, B. B. et al. Toward discovery science of human brain function. Proceedings of the National Academy of Sciences 107, 4734–4739, doi:10.1073/pnas.0911855107 (2010).

23 Schumacher, J. et al. Functional connectivity in dementia with Lewy bodies: A within- and between-network analysis. Human brain mapping 39, 1118–1129, doi:10.1002/hbm.23901 (2018).

24 Kenny, E. R., Blamire, A. M., Firbank, M. J. & O’Brien, J. T. Functional connectivity in cortical regions in dementia with Lewy bodies and Alzheimer’s disease. Brain: a journal of neurology 135, 569–581, doi:10.1093/brain/awr327 (2012).

25 Peraza, L. R. et al. Resting state in Parkinson’s disease dementia and dementia with Lewy bodies: commonalities and differences. International journal of geriatric psychiatry 30, 1135–1146, doi:10.1002/gps.4342 (2015).

26 Schneider, J. A., Arvanitakis, Z., Bang, W. & Bennett, D. A. Mixed brain pathologies account for most dementia cases in community-dwelling older persons. Neurology 69, 2197–2204, doi:10.1212/01.wnl.0000271090.28148.24 (2007).

27 Shimada, H. et al. beta-Amyloid in Lewy body disease is related to Alzheimer’s disease-like atrophy. Movement disorders: official journal of the Movement Disorder Society 28, 169–175, doi:10.1002/mds.25286 (2013).

28 Liao, X., Cao, M., Xia, M. & He, Y. Individual differences and time-varying features of modular brain architecture. NeuroImage 152, 94–107, doi:10.1016/j.neuroimage.2017.02.066 (2017).

29 Craddock, R. C., James, G. A., Holtzheimer, P. E., Hu, X. P. & Mayberg, H. S. A whole brain fMRI atlas generated via spatially constrained spectral clustering. Human brain mapping 33, 1914–1928, doi:10.1002/hbm.21333 (2012).

30 Bruno, R. M. & Sakmann, B. Cortex is driven by weak but synchronously active thalamocortical synapses. Science 312, 1622–1627, doi:10.1126/science.1124593 (2006).

31 Bassett, D. S., Yang, M., Wymbs, N. F. & Grafton, S. T. Learning-induced autonomy of sensorimotor systems. Nature neuroscience 18, 744, doi:10.1038/nn.3993 https://www.nature.com/articles/nn.3993#supplementary-information (2015).

32 Hoshi, E., Tremblay, L., Féger, J., Carras, P. L. & Strick, P. L. The cerebellum communicates with the basal ganglia. Nature neuroscience 8, 1491, doi:10.1038/nn1544 https://www.nature.com/articles/nn1544#supplementary-information (2005).

33 Takahashi, E., Hayashi, E., Schmahmann, J. D. & Ellen Grant, P. Development of cerebellar connectivity in human fetal brains revealed by high angular resolution diffusion tractography. NeuroImage 96, 326–333, doi:https://doi.org/10.1016/j.neuroimage.2014.03.022 (2014).

34 Yeo, B. T. et al. Functional Specialization and Flexibility in Human Association Cortex. Cerebral cortex 25, 3654–3672, doi:10.1093/cercor/bhu217 (2015).

35 Buckner, R. L. & Krienen, F. M. The evolution of distributed association networks in the human brain. Trends in cognitive sciences 17, 648–665, doi:10.1016/j.tics.2013.09.017 (2013).

36 Allman, J. M. et al. The von Economo neurons in frontoinsular and anterior cingulate cortex in great apes and humans. Brain Structure and Function 214, 495–517, doi:10.1007/s00429-010-0254-0 (2010).

37 Menon, V. & Uddin, L. Q. Saliency, switching, attention and control: a network model of insula function. Brain structure & function 214, 655–667, doi:10.1007/s00429-010-0262-0 (2010).

38 Turchi, J. et al. The Basal Forebrain Regulates Global Resting-State fMRI Fluctuations. Neuron 97, 940–952.e944, doi:10.1016/j.neuron.2018.01.032 (2018).

39 Goulas, A., Schaefer, A. & Margulies, D. S. The strength of weak connections in the macaque cortico-cortical network. Brain structure & function 220, 2939–2951, doi:10.1007/s00429-014-0836-3 (2015).

40 Kaiser, M., Hilgetag, C. C. & van Ooyen, A. A simple rule for axon outgrowth and synaptic competition generates realistic connection lengths and filling fractions. Cerebral cortex 19, 3001–3010, doi:10.1093/cercor/bhp071 (2009).

41 Crossley, N. A. et al. Cognitive relevance of the community structure of the human brain functional coactivation network. Proceedings of the National Academy of Sciences 110, 11583–11588, doi:10.1073/pnas.1220826110 (2013).

42 Lemaitre, H. et al. Normal age-related brain morphometric changes: nonuniformity across cortical thickness, surface area and gray matter volume? Neurobiology of aging 33, 617.e611–619, doi:10.1016/j.neurobiolaging.2010.07.013 (2012).

43 Frisoni, G. B., Prestia, A., Rasser, P. E., Bonetti, M. & Thompson, P. M. In vivo mapping of incremental cortical atrophy from incipient to overt Alzheimer’s disease. Journal of neurology 256, 916–924, doi:10.1007/s00415-009-5040-7 (2009).

44 Firbank, M. et al. Neural correlates of attention-executive dysfunction in lewy body dementia and Alzheimer’s disease. Human brain mapping, doi:10.1002/hbm.23100, doi:10.1002/hbm.23100 (2015).

45 Santarnecchi, E., Galli, G., Polizzotto, N. R., Rossi, A. & Rossi, S. Efficiency of weak brain connections support general cognitive functioning. Human brain mapping 35, 4566–4582, doi:10.1002/hbm.22495 (2014).

46 Seeley, W. W., Crawford, R. K., Zhou, J., Miller, B. L. & Greicius, M. D. Neurodegenerative diseases target large-scale human brain networks. Neuron 62, 42–52, doi:10.1016/j.neuron.2009.03.024 (2009).

47 Zhou, J., Gennatas, E. D., Kramer, J. H., Miller, B. L. & Seeley, W. W. Predicting regional neurodegeneration from the healthy brain functional connectome. Neuron 73, 1216–1227, doi:10.1016/j.neuron.2012.03.004 (2012).

48 McKeith, I. G. et al. Diagnosis and management of dementia with Lewy bodies: Fourth consensus report of the DLB Consortium. Neurology 89, 88–100, doi:10.1212/wnl.0000000000004058 (2017).

49 McKeith, I. G. et al. Diagnosis and management of dementia with Lewy bodies: third report of the DLB Consortium. Neurology 65, 1863–1872, doi:10.1212/01.wnl.0000187889.17253.b1 (2005).

50 Emre, M. et al. Clinical diagnostic criteria for dementia associated with Parkinson’s disease. Movement disorders: official journal of the Movement Disorder Society 22, 1689–1707; quiz 1837, doi:10.1002/mds.21507 (2007).

51 McKhann, G. M. et al. The diagnosis of dementia due to Alzheimer’s disease: Recommendations from the National Institute on Aging-Alzheimer’s Association workgroups on diagnostic guidelines for Alzheimer’s disease. Alzheimer’s & dementia: the journal of the Alzheimer’s Association 7, 263–269, doi:10.1016/j.jalz.2011.03.005 (2011).

52 Brown, J., Rudie, J., Bandrowski, A., Van Horn, J. & Bookheimer, S. The UCLA multimodal connectivity database: a web-based platform for brain connectivity matrix sharing and analysis. Frontiers in neuroinformatics 6, doi:10.3389/fninf.2012.00028 (2012).

53 Jenkinson, M., Beckmann, C. F., Behrens, T. E., Woolrich, M. W. & Smith, S. M. Fsl. NeuroImage 62, 782–790 (2012).

54 Patel, A. X. et al. A wavelet method for modeling and despiking motion artifacts from resting-state fMRI time series. NeuroImage 95, 287–304, doi:http://dx.doi.org/10.1016/j.neuroimage.2014.03.012 (2014).

55 Peraza, L. R. et al. Intra- and inter-network functional alterations in Parkinson’s disease with mild cognitive impairment. Human brain mapping, doi:10.1002/hbm.23499 (2017).

56 Baggio, H.-C. et al. Functional brain networks and cognitive deficits in Parkinson’s disease. Human brain mapping 35, 4620–4634, doi:10.1002/hbm.22499 (2014).

57 Schwarz, A. J. & McGonigle, J. Negative edges and soft thresholding in complex network analysis of resting state functional connectivity data. NeuroImage 55, 1132–1146, doi:10.1016/j.neuroimage.2010.12.047 (2011).

58 Keller, C. J. et al. Neurophysiological Investigation of Spontaneous Correlated and Anticorrelated Fluctuations of the BOLD Signal. The Journal of Neuroscience 33, 6333–6342, doi:10.1523/jneurosci.4837-12.2013 (2013).

59 Schafer, K. et al. Negative BOLD signal changes in ipsilateral primary somatosensory cortex are associated with perfusion decreases and behavioral evidence for functional inhibition. NeuroImage 59, 3119–3127, doi:10.1016/j.neuroimage.2011.11.085 (2012).

60 Kelly, A. M. C., Uddin, L. Q., Biswal, B. B., Castellanos, F. X. & Milham, M. P. Competition between functional brain networks mediates behavioral variability. NeuroImage 39, 527–537, doi:http://dx.doi.org/10.1016/j.neuroimage.2007.08.008 (2008).

61 Achard, S. & Bullmore, E. Efficiency and Cost of Economical Brain Functional Networks. PLoS computational biology 3, e17, doi:10.1371/journal.pcbi.0030017 (2007).

62 Brito, M. R., Chávez, E. L., Quiroz, A. J. & Yukich, J. E. Connectivity of the mutual k-nearest-neighbor graph in clustering and outlier detection. Statistics & Probability Letters 35, 33–42, doi:https://doi.org/10.1016/S0167-7152(96)00213-1 (1997).

63 Jalili, M. Functional Brain Networks: Does the Choice of Dependency Estimator and Binarization Method Matter? Scientific reports 6, 29780, doi:10.1038/srep29780 (2016).

64 Newman, M. Finding community structure in networks using the eigenvectors of matrices. Physical Review E 74, 036104 (2006).

65 Mammen, E. in When Does Bootstrap Work? 106–117 (Springer, 1992).

66 Mandke, K. et al. Comparing multilayer brain networks between groups: Introducing graph metrics and recommendations. NeuroImage 166, 371–384, doi:https://doi.org/10.1016/j.neuroimage.2017.11.016 (2018).

67 Steen, M., Hayasaka, S., Joyce, K. & Laurienti, P. Assessing the consistency of community structure in complex networks. Physical Review. E, Statistical, Nonlinear, and Soft Matter Physics 84, 016111–016111 (2011).

68 Doron, K. W., Bassett, D. S. & Gazzaniga, M. S. Dynamic network structure of interhemispheric coordination. Proceedings of the National Academy of Sciences of the United States of America 109, 18661–18668, doi:10.1073/pnas.1216402109 (2012).

69 Xia, M., Wang, J. & He, Y. BrainNet Viewer: A Network Visualization Tool for Human Brain Connectomics. PloS one 8, e68910, doi:10.1371/journal.pone.0068910 (2013).

70 Moreland, K. 92–103 (Springer Berlin Heidelberg).

